# Avian germline-restricted chromosomes are reservoirs for active long-terminal-repeat retroviruses

**DOI:** 10.1101/2025.06.27.662071

**Authors:** Bohao Fang, Scott V. Edwards

## Abstract

Germline-restricted chromosomes (GRCs) are unique to germ cells and absent from somatic cells in songbirds. However, their contents, functions, and evolutionary mechanisms remain unclear. We performed comparative genomics on long-read assembled GRCs from male House Finch (*Haemorhous mexicanus*), Common Rosefinch (*Carpodacus erythrinus*), and Blue Tit (*Cyanistes caeruleus*), the first two of which are newly presented and annotated here. These long-read GRCs are repeat-rich (63–74%), despite their wide variation in size (16–160 megabases), individually distinct gene content, and variable evolutionary histories. These GRCs have accumulated intact long terminal repeat (LTR) endogenous retroviruses (ERVs), and their proliferation co-occurs with speciation events of passerine birds. In contrast, normal chromosomes (autosomes and sex chromosomes) are known to purge ERVs through ectopic recombination of LTRs that generate solo LTRs. Furthermore, transcriptomic data reveal significantly higher ERV transcription on GRCs than on A-chrs in House Finch testis. We propose that GRCs act as reservoirs of active ERVs, promoting germline transposition and potentially accelerating adaptation and speciation. This study suggests that GRCs play an indispensable role and likely contribute to adaptive diversification in birds.

**Graphical abstract:** 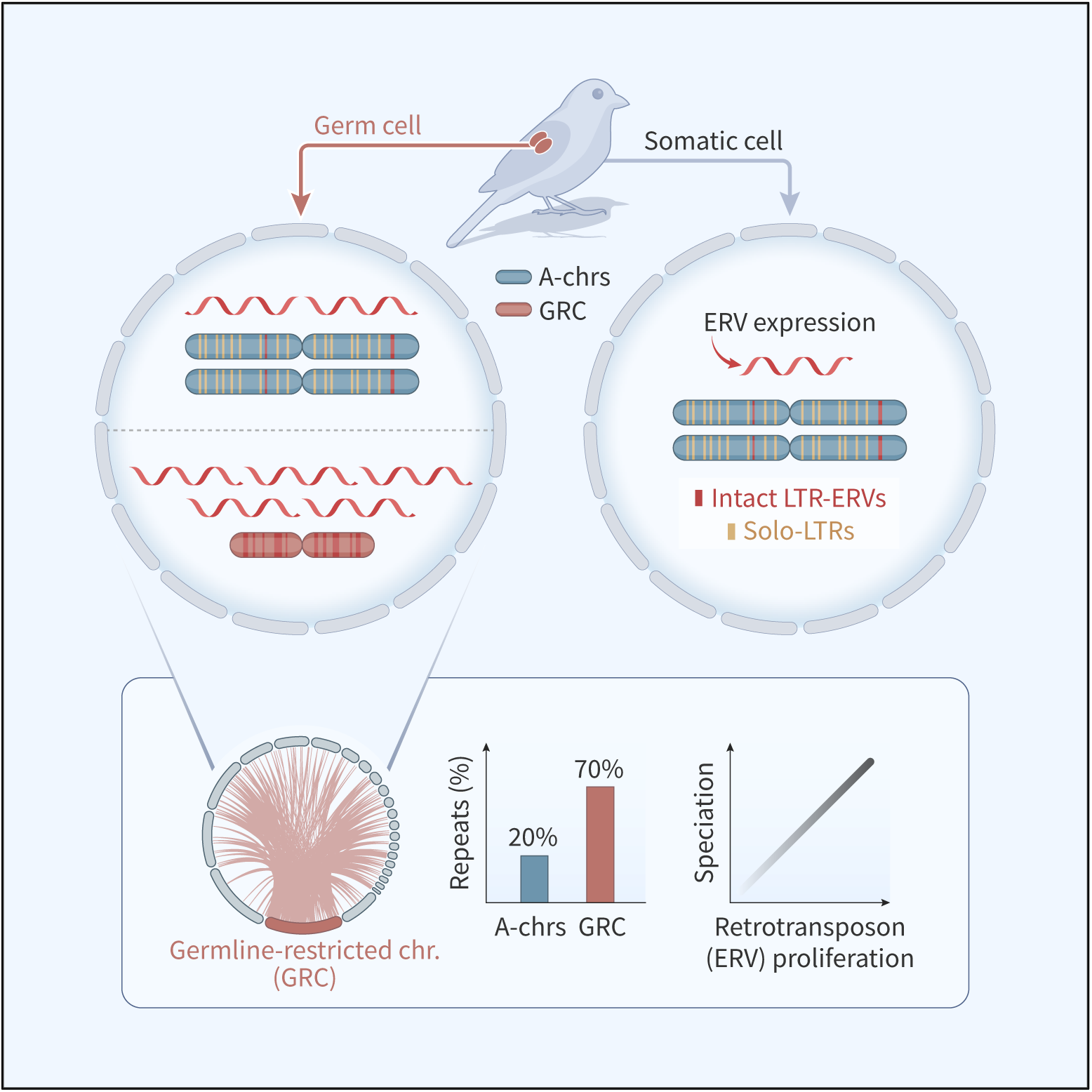

## Introduction

The discovery of germline-restricted chromosomes (GRCs) in songbirds has reshaped our understanding of avian genome organization ^1^. GRCs are present in the germ cells (spermatocytes and oocytes) but are eliminated from somatic cells in numerous Passeriformes, the largest avian order ^2^. First documented in the zebra finch (*Taeniopygia guttata*) (Pigozzi & Solari 1998), GRCs have since been detected in every songbird examined—that is, in oscine passerines (suborder *Passeri*, the largest radiation within Passeriformes)—showing unusual inheritance patterns and ranging in size from micro-to macro-chromosomes, which underscores their rapid evolution across species ^3^. Most research on GRCs to date has focused on their cytogenetic characteristics ^1,4^. However, despite their ubiquity, base-level sequencing data have lagged behind. There are only five published GRCs with sequencing data ^4^, four of which are based on short-read sequencing, and thus fundamental questions remain about their origin, contents, functions, and evolutionary mechanisms ^5^.

Transposable elements (TEs), and endogenous retroviruses (ERVs) in particular, have emerged as key drivers of eukaryotic genome evolution and speciation through their capacity to shape gene regulatory networks, create structural rearrangements, and in some cases promote reproductive barriers ^6–10^. Bursts of TEs have been correlated with diversification in multiple vertebrate clades. For example, mammals present a strong positive correlation between transposon insertion density and speciation rate across mammalian families ^11^. Similar associations have been documented for primates ^12^, bats ^13^ and Anolis lizards ^14^. Avian genomes were widely regarded as streamlined and repeat-poor compared to other taxa ^15^. However, growing evidence suggests that avian lineages have undergone significant TE activity, especially on the female-specific W chromosome ^16^. A recent study also proposes that solo-LTR insertions can act as regulatory elements, fueling phenotypic diversification, underscoring the evolutionary significance of TE-mediated mechanisms in birds ^17^.

Early studies noted the presence of multiple gene duplicates on avian GRCs, many of which are implicated in germline or early developmental processes ^18^. However, unlike many other well-known germline-limited systems (hagfish, ciliates, and lamprey) whose GRCs accumulate large amounts of repetitive DNA because of limited recombination ^19,20^, the repeat content on avian GRCs is still poorly characterized.

Here, we use PacBio highly accurate long-read (HiFi) sequencing of germline and somatic tissues from the House Finch (*Haemorhous mexicanus*) and Common Rosefinch (*Carpodacus erythrinus*), integrating data from the only other published long-read-assembled GRC from the Blue Tit (*Cyanistes caeruleus*) ^21^. The House Finch is native to North America, whereas the Common Rosefinch spans Eurasia; molecular phylogenies place their split within Fringillidae at ∼12 Ma ^22^. By assembling and annotating these GRCs, we aim to (i) identify GRC-linked genes and their evolutionary trajectories, (ii) assess their repeat content, and (iii) test whether the magnitude and timing of repeat accumulation on GRCs track major pulses of passerine diversification.

## Results

### Sequencing and Assembling GRCs

To assemble GRC and normal chromosomes (autosomes and sex chromosomes, collectively called ‘A-chromosomes [A-chrs]’) of House Finch and Common Rosefinch, we obtained HiFi sequencing data from germline (testis) and somatic (heart) tissues of the same male individuals for each species. For comparative analyses, we also incorporated previously published HiFi-based assemblies of A-chrs and GRC of a male Blue Tit ^21^, providing a third reference point for interspecies comparisons. Because avian testis comprise both germ and somatic cells, all ‘testis’ libraries inherently contain a mixture of germline and somatic reads. High-depth HiFi reads were obtained for heart and testis in House Finch (HF) and Rosefinch (RF)—HF: heart 33.3×, testis 53.4×; RF: heart 45.6×, testis 98.5× (Table S1)—with HF testis and RF heart newly sequenced here, and HF heart and RF testis coverage augmented using HiFi reads from Fang and Edwards ^23^ (Methods). The Rosefinch A-chrs were assembled directly from somatic HiFi reads using hifiasm ^24^ (Methods), resulting in an assembly of 1.2 gigabases (Gb) in size (582 contigs; N50=25.6 megabytes [Mb], BUSCO= 95.5%; Table S2). The existing House Finch chromosome-level somatic assembly ^23^ (39 autosomes plus Z, W sex chromosomes; N50=77.2 Mb) was used as its A-chrs in this study.

To assemble the GRCs, we used a four-step HiFi approach modified from Mueller *et al.* ^21^ and Schlebusch et al. ^25^ (Fig. S1): (1) we mapped somatic reads to the assembled germline contigs (A-chrs plus GRC) (2) identified GRC-specific regions by searching for intervals in the germline assembly with zero somatic read coverage, (3) extracted germline reads mapping to those putative GRC regions, and (4) re-assembled these extracted reads *de novo* (see Methods for details). Given the more extensive genomic resources available for House Finch ^23^, additional short-read whole genome sequencing with additional individuals (21–33×) helped refine the House Finch GRC assembly. These additional analyses likely contribute to a more complete GRC assembly for House Finch than for Rosefinch. However, using these four-steps, most of the Rosefinch GRC should be captured, as long as its sequences are divergent from A-chrs ^21^.

Both assembled GRCs are large chromosomes (macro-GRCs) (Fig. 1). The House Finch GRC is 168.4 Mb, composed of 947 contigs (N50=0.79 Mb; Table S2), and is the largest chromosome in the House Finch genome. The Rosefinch GRC is 54.49 Mb (1,002 contigs; N50=75,259 base pairs [bp]; Table S2). We validated the assembled GRCs by mapping somatic and germline HiFi reads to the combined genome (A-chr plus GRC; Fig. S2) and checking their coverage: (1) the GRC contigs showed zero coverage from somatic reads, indicating that GRC contigs were unique to germline tissue; (2) in testis, GRC coverage was lower than A-chr coverage (House Finch: 102.32 vs. 21.98; Rosefinch: 99.77 vs. 29.35), consistent with the GRC’s haploid status and presence only in spermatocytes, as also observed in the Blue Tit GRC ^21^; and (3) the “GRC / A-chr” coverage ratios (House Finch = 0.22, Rosefinch = 0.29) closely matched the ratios reported in other songbird testis-based GRC sequencing studies ^5,21,25^. We note, however, that many gaps in our GRC assembly remain despite using HiFi sequencing, meaning our estimates of GRC length is conservative.

**Figure 1.**
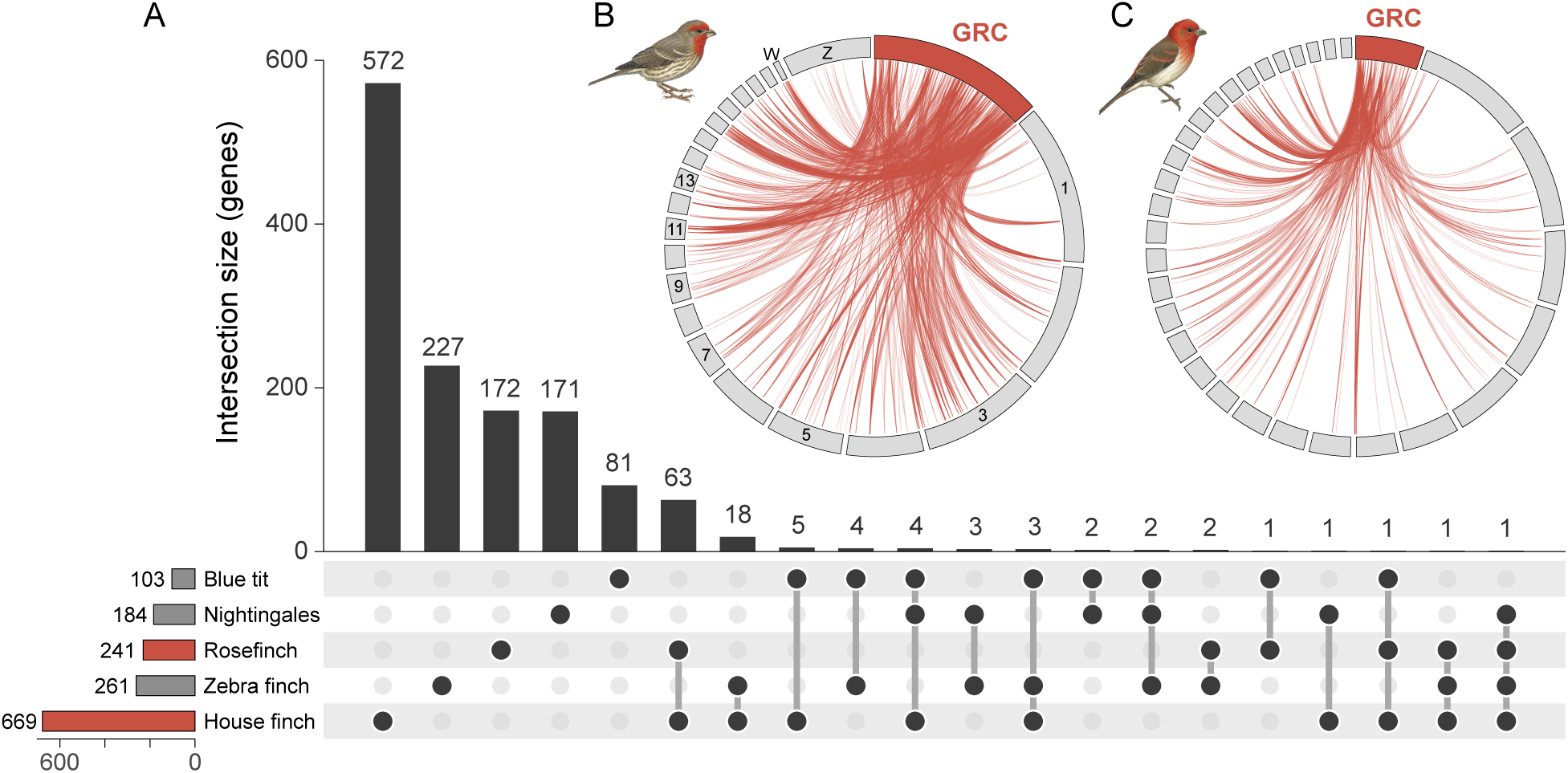
Assembled germline-restricted chromosomes (GRCs) and shared GRC-linked genes across bird species. (A) Intersection size of GRC-linked genes shared among different bird species with published GRCs. The vertical bar chart represents the number of shared genes, and the horizontal bars indicate the total number of GRC genes annotated per species. (B, C) Circos plot showing GRC-linked genes and their A-chromosomal paralogs in the House Finch; B) and the Common Rosefinch (C). The largest 20 autosomes and sex chromosomes are displayed in (B), and scaffolds larger than 100 Mb are displayed in (C).

### Functionally Important Genes in Avian GRCs

To annotate protein-coding genes on the newly assembled GRCs and A-chrs, we performed *ab initio* and evidence-driven gene predictions informed by RNA-seq data (from gonad, brain, and heart tissues). We curated gene models for completeness, assessed homology by blastx ^26^ against NCBI’s protein database, and used the RNA-seq data to quantify gene expression across GRCs and A-chrs (Methods). This approach identified and annotated 669 genes on the House Finch GRC and 241 in the Rosefinch GRC, compared to 20,054 in House Finch A-chrs and 14,490 in Rosefinch A-chrs (Table S2). Notably, the House Finch GRC includes paralogs of genes originating from all 39 House Finch autosomes and both Z and W sex chromosomes, including 13 genes from the W chromosome (W-chr; Fig. 1), eight of which exhibit detectable expression (Fig. S3). These data imply that gene content on the male GRC is derived not only from autosomes but also from the female W-chr, thus supporting the hypothesis that the GRC may act as a reservoir for potentially conflicting genes, thereby reducing sexual conflict by balancing gene dosage between sexes ^27^. Although the Rosefinch A-scaffolds are not assigned to chromosomes, GRC-linked genes in this species contain paralogs from all scaffolds larger than 100 Mb (Fig. 1).

GRCs harbor both functional genes and pseudogenes in high copy numbers: 387 (57.8%) genes in House Finch and 102 (42.3%) genes in Rosefinch exhibit duplications within the GRC. NANO3 in House Finch GRC is noteworthy as one of only two ‘universal’ genes in germ cell development, functioning across all animals so far investigated, from flies, worms, frogs, mice to humans ^28^.

NANO3 exhibited the highest number of gene copies on the GRC and a single copy in House Finch A-chrs; however, of 286 copies only one was expressed on the GRC (Fig. S4). Summarizing published GRCs of songbirds, including the Common and Thrush Nightingales (*Luscinia megarhynchos* and *L. luscinia*) and the Zebra Finch (*Taeniopygia castanotis*), we observe that the majority (91.7%; n=1,223) of GRC-linked genes are species-specific (Fig. 1A). Yet 12 GRC-linked genes (e.g., CPEB1, associated with germline development) are shared in more than two species (Fig. 1), suggesting an essential function. Gene ontology (GO) analyses suggest that many GRC-linked genes function in gametogenesis, cell-cycle control, or early embryo development (Fig. S5), suggesting they are key players in germline development.

### Ancient Retention and Recent Turnover of GRC-Linked Genes

To infer their evolutionary origins, 12 GRC-linked genes shared in more than two species were used for phylogenetic inference across major Passeriformes lineages. To span the major lineages of Passeriformes, we retrieved orthologous coding sequences from Chicken (*Gallus gallus*), the Rifleman (*Acanthisitta chloris*), and two Suboscines (Blue-capped Manakin, *Lepidothrix coronata*, and Golden-collared Manakin, *Manacus vitellinus*), and constructed phylogenies and performed an approximately unbiased (AU) topology test ^29^ (Methods). The results reveal variable evolutionary histories of these GRC-linked genes: Two of the genes (CPEB1 and ELAVL4) evolved before the most recent common ancestor of oscines and suboscines (scenario I); five genes (scenario II) duplicated before the diversification of oscines; and 5 genes duplicated within the oscines (scenario III; Fig. 2A; Fig. S6). The most broadly retained gene, CPEB1, present in 4/5 studied GRCs (Fig. 1A), may trace back to an ancestral Passeriformes lineage before the divergence of oscines and suboscines (∼50 million years ago [Mya], Fig. 2D). These findings suggest that although most GRC-linked genes are species-specific and recently arisen within oscines (songbirds), a subset predates the oscine-suboscine split and thus may have existed throughout passerine evolution.

**Figure 2.**
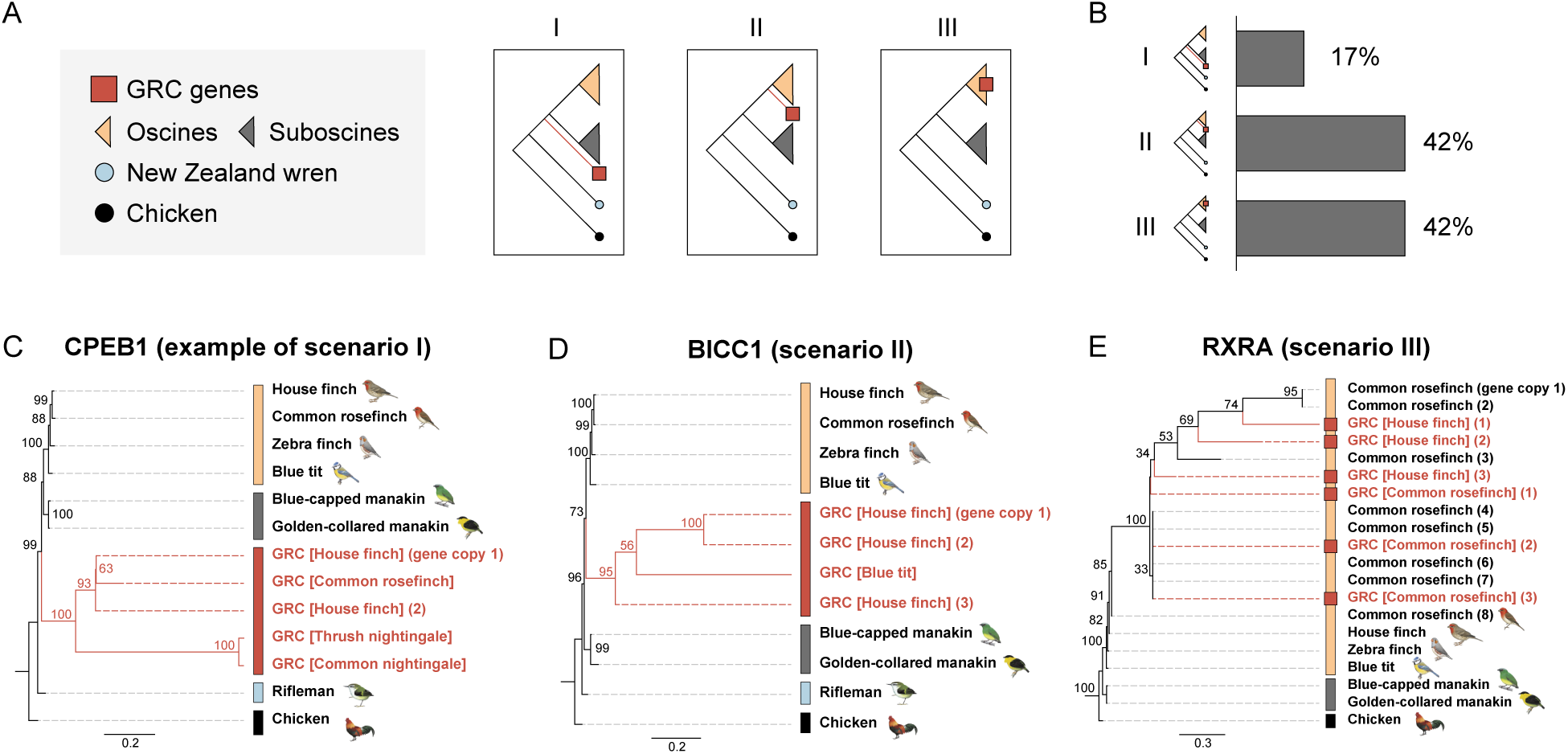
Dynamic evolutionary histories of GRC-linked genes shared across bird species. (A) Schematics of hypothesized scenarios of evolutionary histories of GRC-linked genes: scenario I (evolving before the divergence of both Oscines and Suboscines), scenario II (GRC genes evolved before the divergence of Oscines), and scenario III (evolving within Oscines). (B) Percentage of GRC-genes fitting each evolutionary scenario (I, II, III) among 12 GRC-genes shared across at least three bird species. (C–E) Examples of phylogenetic trees of specific genes under each scenario: (C) CPEB1 (I), (D) BICC1 (II), and (E) RXRA (III). Phylogenetic trees were constructed with 1,000 bootstrap replicates, and topological robustness was evaluated with an approximately unbiased test.

### Enrichment of Repeats and Transposable Elements (TEs) on Avian GRCs

We next annotated repetitive elements in both GRCs and A-chrs for House Finch, Rosefinch, and Blue Tit using RepeatMasker ^30^ and LTR_retriever ^31^ (Methods). The latter program integrates outputs from LTR_harvest ^32^ and LTR_FINDER ^33^ to produce high-quality, non-redundant LTR libraries and annotations and has been used to identify long terminal repeat retrotransposons (LTR-RTs) in diverse clades (citations). The results reveal a consistently high abundance of repeats across GRCs (House Finch: 72.6%, Rosefinch: 73.6% and Blue Tit: 63.2%), in contrast to lower abundance across A-chrs (17.9%, 20.8% and 20.8%; Table S4; Fig. 3A–C). LTR and LINE (long interspersed nuclear elements) are consistently the two major repeat types in A-chrs and GRCs, having begun to diversify 50–60 Mya (Fig. 3), about the time of the most recent common ancestor of passerine birds ^34,35^ (Fig. 3D–I; Methods). GRCs contain a substantially higher proportion of transposable elements compared to A-chrs, particularly LTRs, which comprise 23.0–40.3% of GRCs compared to only 5.8– 7.2% of A-chrs. Endogenous retroviruses (ERVs) are the predominant family of LTRs, constituting over 90% of classified LTRs across long-read GRCs (Fig. S6), suggesting that GRCs have experienced extensive integration of retroviral elements during their evolution.

**Figure 3.**
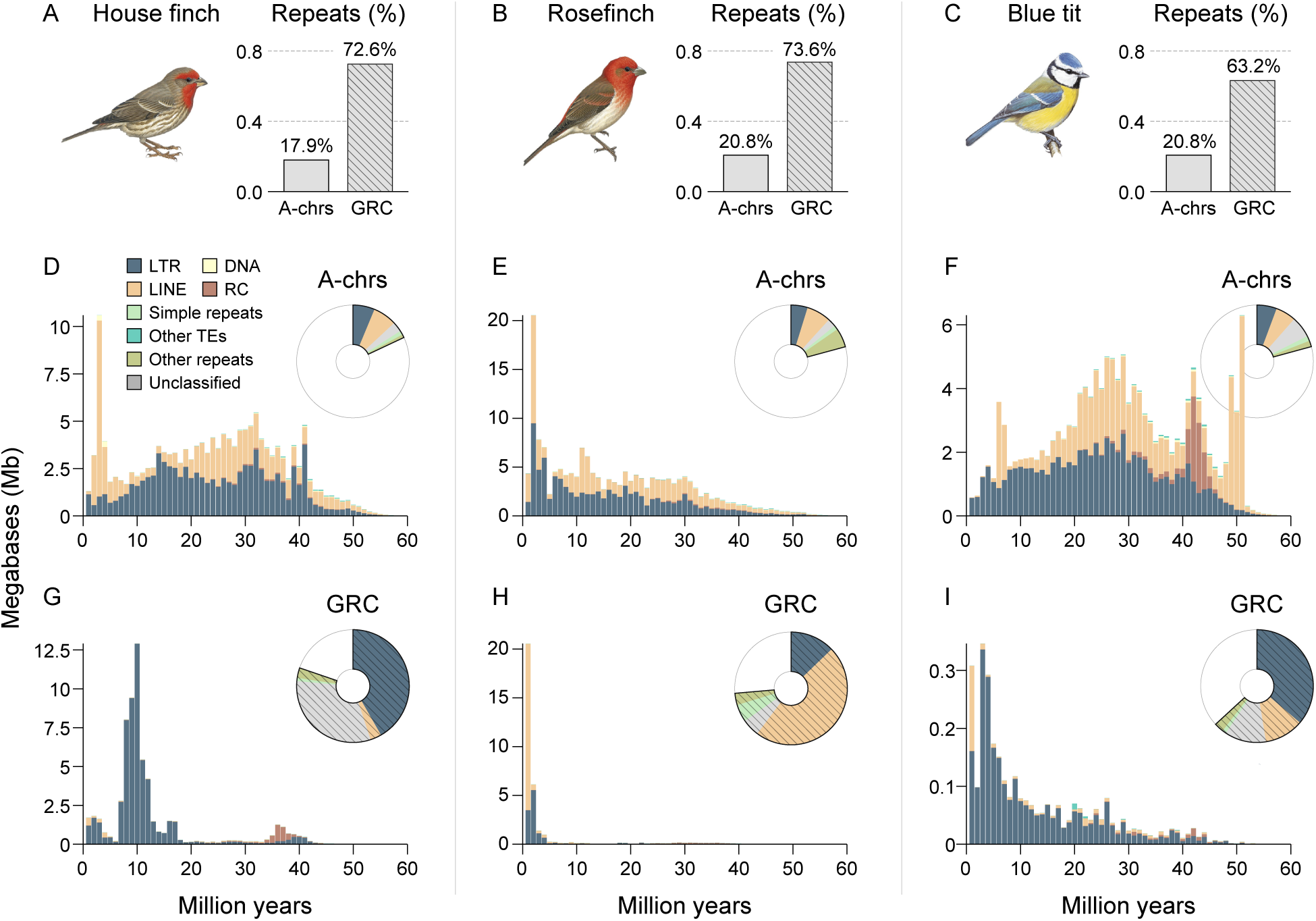
Avian GRC is repeat-rich. (A–C) The bar charts show the percentage of total repeat content in A-chromosomes (A-chrs) and GRCs of (A) House Finch, (B) Common Rosefinch, and (C) Blue Tit, whose GRCs were assembled using long-read sequencing. (D–I) Temporal landscape of transposable elements (TEs) in A-chrs (D–F) and GRCs (G–I) of the three species. The TE landscapes are displayed as stacked histograms in bins of 1 million years (My), using a substitution rate of 3.3×10^-3^ substitutions per site per million years for Passeriformes (referencing Zhang et al., 2014; Methods). The pie charts show the composition of repeats. LINE, long interspersed nuclear elements; RC, Rolling Circle.

### Endogenous Retrovirus (ERVs) are Retained on GRCs but Purged from A-chrs

LTR retrotransposons enter the genome as full-length elements bounded by two identical long terminal repeats (intact LTRs). Solo-LTRs are typically generated by ectopic recombination between flanking LTR element ^36,37^ and can function as regulatory elements ^38,39^, potentially contributing to the adaptive diversification of passerine birds (Chen et al. 2024a) and other vertebrates ^39,40^. We identified intact LTRs and solo-LTRs from both GRCs and A-chrs in the three species using Perl scripts in LTR_retriever under stringent criteria (Methods). To quantify recombination-mediated deletion of ERVs on these chromosomes, we computed the ratio of solo to intact LTRs for each LTR family. Our results confirm that A-chrs harbor high solo/intact ratios: House Finch A-chrs average 80.3 (median 17.3; max 954), Common Rosefinch A-chrs 32.7 (median 8.5; max 477), and Blue Tit A-chrs 39.4 (median 7.5; max 798) (Fig. 4). This observation aligns with previous findings in avian A-chrs, where the majority of ERV copies are present as solo-LTRs while intact LTRs are rare ^41^. By contrast, GRCs show greatly reduced ratios—10.8 (median 3.0; max 46.3) in House Finch, 1.41 (median 1.0; max 4.0) in Rosefinch, and 3.99 (median 3.5; max 10.0) in Blue Tit—representing 7.4-, 23.2-, and 9.9-fold lower values than their corresponding A-chrs, respectively (Fig. 4). These data indicate preferential retention on GRCs of LTR pairs flanking ERV coding regions that are essential for retro-transposition.

**Figure 4.**
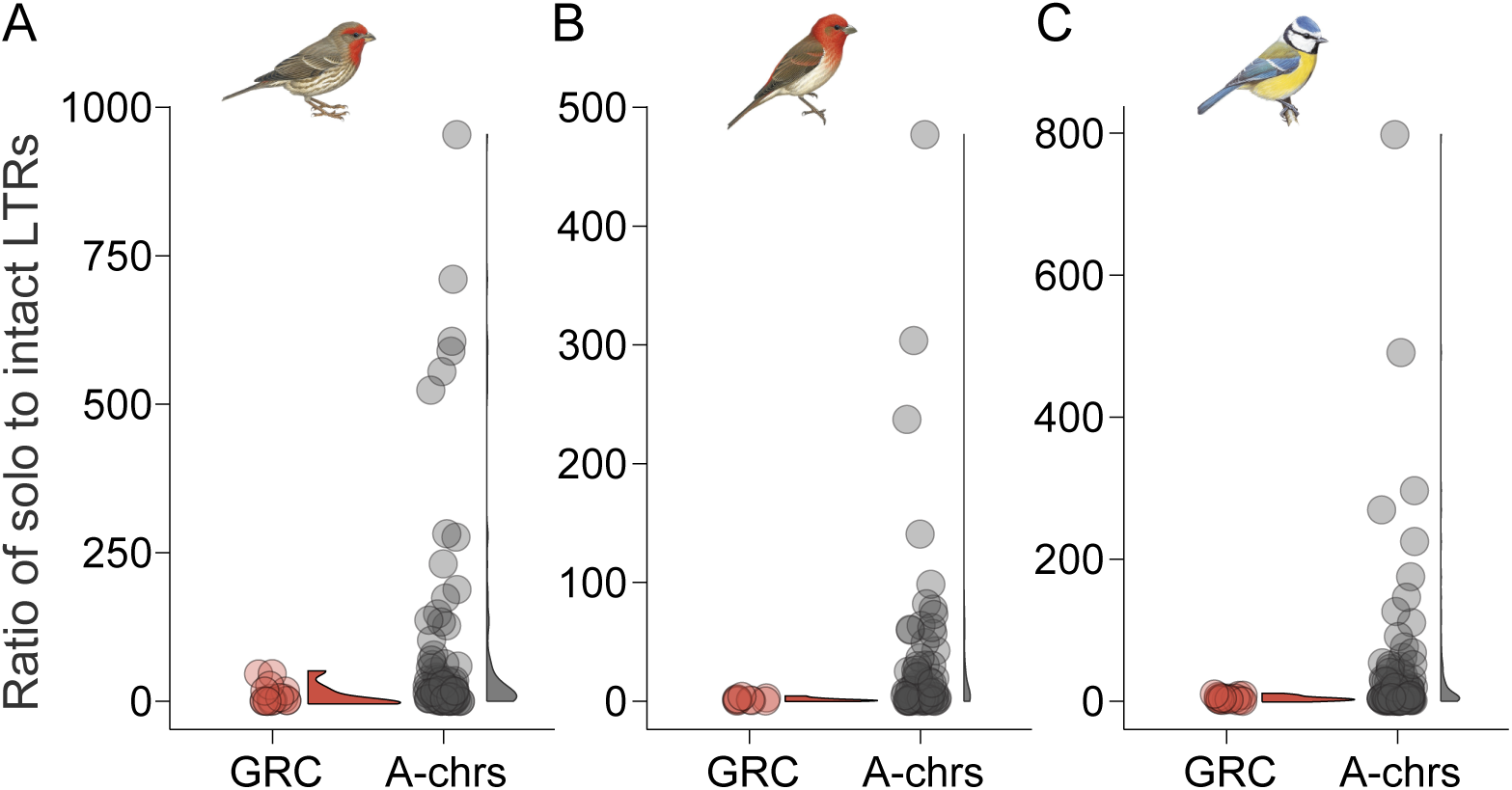
Proliferation of Solo-LTRs on A-chromosomes and retention of intact-LTR on GRCs. (A–C) Ratio of ERV-derived solo-LTRs to intact-LTRs in GRCs and A-chrs of (A) House Finch, (B) Common Rosefinch, and (C) Blue Tit. Solo LTRs are formed typically by ectopic recombination between two LTR regions at non-homologous locations on the same or different chromosomes. The solo-intact LTR ratio was calculated by dividing the number of solo LTRs by the number of intact LTR-RTs for each LTR family.

### Active Transcription of Endogenous Retroviral (ERV) Genes in GRCs

Considering the exceptionally high number of GRC-linked intact ERV-LTRs, we tested whether these ERVs are active in germline tissue (testis) and within the GRC in House Finches, using ERV gene transcription. Paired-end RNA-seq reads were trimmed and mapped to the respective assemblies: brain and heart reads were aligned to the somatic assembly (A-chrs), whereas testis reads were aligned to the germline assembly (A-chrs plus GRC). We then quantified RNA expression levels for ERV retroviral genes ^42^ and evaluated differential expression between gonadal and somatic tissues, normalized for sequencing depth and gene length, as well as between A-chrs and GRC in germline cells, normalized based on the estimated fraction of germline cells present in testis library, allowing fair comparison between GRCs and A-chrs (Methods). Empirical studies indicate that germline cells constitute a small proportion (commonly less than 5%) of avian testis (e.g., ^43–45^). For instance, Naito *et al.* ^43^ observed that gonocytes represented <1% of total gonadal cells in late-stage chicken embryos, whereas Jung *et al.* ^44^ found that spermatogonial stem cells comprised ∼0.3–0.4% of testicular cells in juvenile chickens and only 0.03–0.07% in adults. Similarly, Hu *et al.* ^45^ reported ∼2– 3% germ cells in embryonic chicken gonads, with subsequent attrition at later developmental stages. We therefore assumed hypothetical germline cell proportions of 10%, 5%, and 1% for the testis sample, covering the above empirical range and considering ∼1% as likely closest to the true value.

Overall, ERV expression was significantly elevated in gonadal tissue compared to somatic tissues. The mean transcript per million (TPM) in testis (412.4) was approximately 1.7 times higher than in brain (237.9) and 1.6 times higher than in heart (255.2) (Fig. 5A, B). This pattern is consistent with observations of greater ERV expression in reproductive tissues (ovary, testis, and primordial germ cells) than in somatic tissues of the zebra finch ^17^. Within testis, ERVs were also expressed at markedly higher levels in GRCs than in A-chrs, even assuming germline cells comprised as much as 10% of the testis tissue (Fig. 5C). Under the assumption of 1% germline cells, ERV expression was 16.6 times higher on GRCs than on A-chrs (mean TPMs: 3570.1 vs. 215.5; Fig. 5C). These findings suggest that intact ERVs retained on avian GRCs are transcriptionally active and capable of germline-specific retrotransposition, potentially also contributing to the overall elevated ERV expression observed in avian gonad tissue.

**Figure 5.**
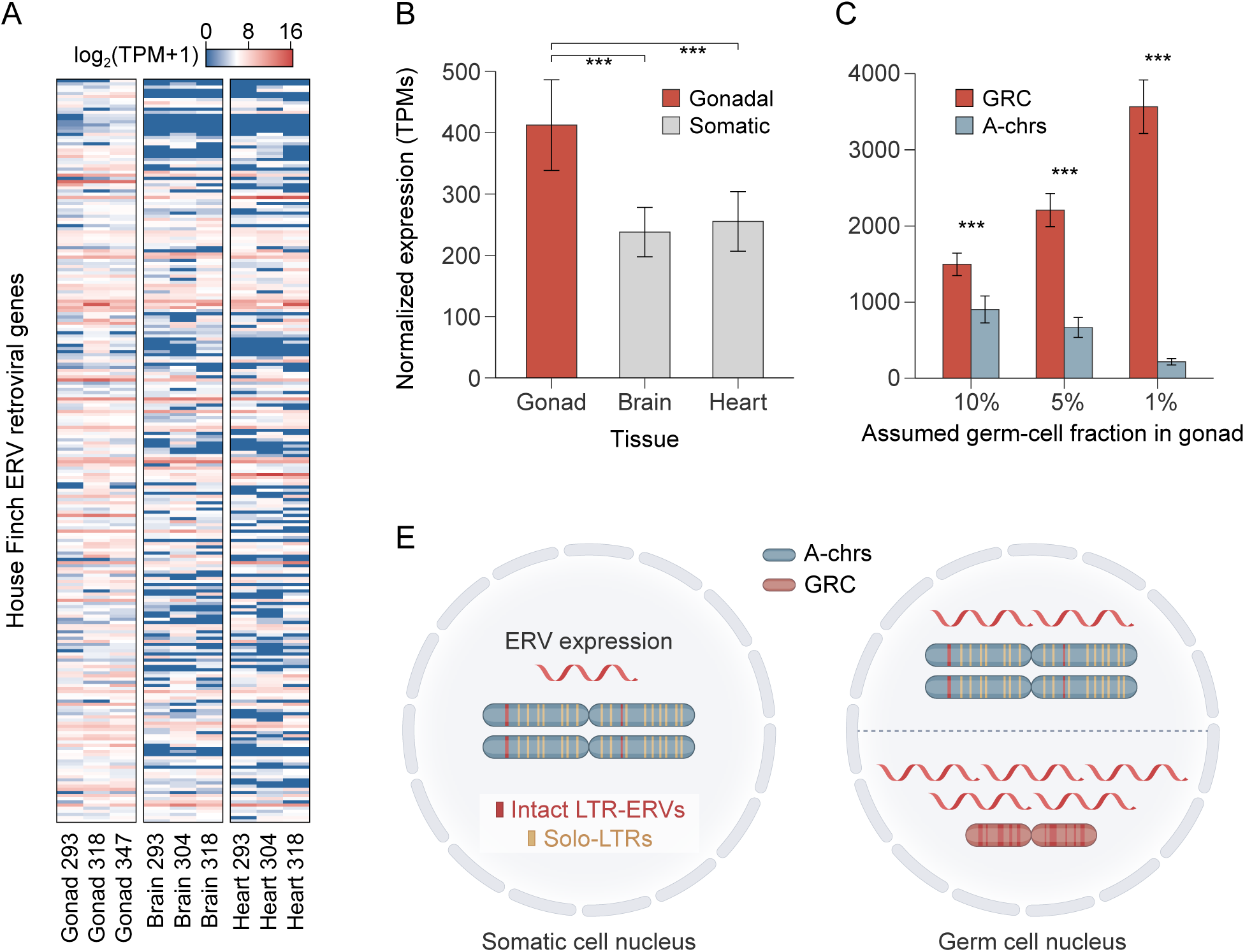
Active ERV retroviral gene transcription in House Finch gonad and GRC. **(A)** Heatmap displaying normalized RNA expression levels (log_2_ (TPM+1)) of ERV retroviral genes across gonad (testis), brain, and heart tissues from four House Finch individuals. Expression is normalized for ERV gene length and sequencing library size. **(B)** Mean normalized expression levels (Transcripts Per Million, TPM ± standard error) of ERV retroviral genes in the three tissues, showing significantly higher expression in gonad compared to somatic tissues (*** p < 0.001, Wald test). **(C)** Comparison of expression levels of ERV retroviral genes between GRCs and A-chrs within gonad tissue. Expression is normalized for gene length, library size, and adjusted assuming hypothetical proportions of germline cells (10%, 5%, 1%) within the testis sample, consistently showing significantly higher expression on GRCs (Wald test). The 1% assumption likely approximates the true biological proportion. **(E)** Schematic model contrasting ERV status and expression in somatic versus germ cell nuclei. A-chrs in both cell types predominantly feature solo-LTRs resulting from ERV purging, exhibiting low transcriptional activity. In contrast, the GRC, unique to germ cells, serves as a reservoir rich in intact LTR-ERVs that show high transcriptional activity, indicating its role in harboring active retroelements.

### Temporal concordance between speciation and bursts of ERV transposition in avian GRCs

We investigated the expansion history of ERVs in both GRCs and A-chrs by profiling the temporal landscapes of ERV long terminal repeats (ERV-LTRs). Our analysis reveals that proliferation of ERV-LTRs through both duplication and insertion follows a similar pattern across the three long-read GRCs, with continuous activity beginning approximately 50 million years ago (Mya), coinciding with the most recent common ancestor of Passeriformes ^34,35^ (Fig. 6). This proliferation peaked at ∼10 Mya (the origin of Passerida, a diverse clade within passerine birds ^2^) and continues to the present. Furthermore, the rate of ERV-LTR proliferation in GRCs, expressed in Mb per million years (Mb/My), is positively correlated with avian speciation events within the TMRCA of major Passeriformes lineages, including Passerida, Muscicapida, Corvides, Passeri (Oscines), and Tyranni (Suboscines). Here, speciation was quantified by counting the number of nodes along the evolutionary path from the common ancestor of Passeriformes to the tips of each species ^17^. In contrast, A-chrs do not show a continuous growth trend of ERV-LTR proliferation over a comparable period of time; indeed, the rate of proliferation of ERV-LTRs on A-chrs is either negatively correlated with (Fig. 6A) or not significantly correlated with (Fig. 6B,C) speciation events. Overall, these results suggest the retention of ERVs and intact LTRs in GRCs during speciation, in contrast to solo-LTR formation via ectopic recombination leading to ERV removal in A-chrs.

**Figure 6.**
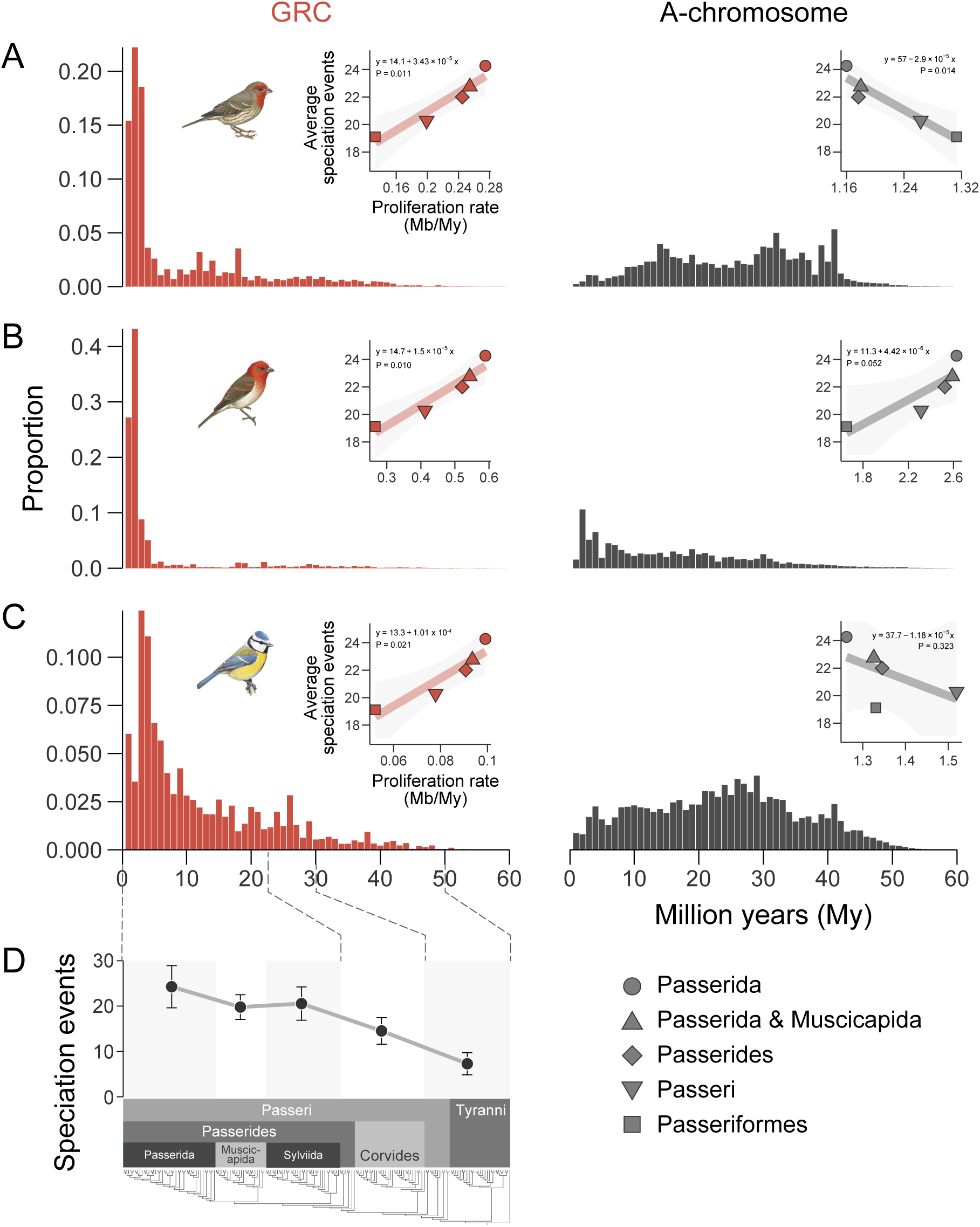
Proliferation of endogenous retroviruses (ERV) in GRCs during Passeriformes speciation. (A–C) Temporal landscape of ERV-LTRs in GRC (left panels) and A-chrs (right panels) for (A) House Finch, (B) Common Rosefinch and (C) Blue Tit. ERV-LTRs in the chromosomes are shown as stacked histograms in 1-million-year (My) bins, using a substitution rate of 3.3×10^-3^ substitutions per site per million years for Passeriformes. Insets show linear regressions illustrating the correlation between ERV-LTR proliferation rates (in Mb/My) and average avian speciation events (see below) within major Passeriformes lineages. (D) Speciation event frequencies across 362 Passeriformes bird species, quantified by node counts from the ancestral Passeriformes node to species tips, based on a family-level phylogeny (modified from Chen et al., 2024). ERV proliferation patterns in GRCs (A–C) show temporal alignment with avian diversification periods (D).

## Discussion

### GRC genomic architecture revealed by long-read sequencing

This study reveals new insights into the biology and evolution of the enigmatic GRCs in songbirds. Our analyses underscore the dynamic evolutionary history of the avian GRC, as evidenced by its substantial variation in size and gene content across species. Despite this variability, we found an exceptionally high density of repeats (63–74%) on GRCs, which are particularly enriched in intact LTR-ERVs, a feature not previously reported for avian GRCs. This repetitive richness stands in stark contrast to most A-chrs, where ectopic recombination purges intact ERVs and leaves solo LTRs— although a few micro-autosomes are themselves unusually repeat-rich ^41^. Furthermore, our transcriptomic data from House Finch testis demonstrate significantly elevated transcription of ERVs on the GRC compared to A-chrs, revealing active transcription of ERVs in the germline.

### Rapid turnover and deeply conserved genes on the GRC

Our annotation illuminates the complex nature of GRC gene content and its evolutionary trajectory. We confirmed previous findings that GRCs harbor numerous gene duplicates and pseudogenes, often having functions in the germline or early development. For example, the House Finch GRC carries hundreds of tandem copies of the germline-determinant gene NANO3, highlighting the chromosome’s bias toward duplicating gametogenesis-related loci ^4^. Our work highlights a dichotomy: the majority of GRC-linked genes appear species-specific, consistent with rapid turnover ^25^, or multiple separate origins of GRCs in passerine birds. However, a core set of genes, such as CPEB1, is conserved across diverse songbirds, suggesting essential, ancestral functions potentially dating to before the oscine-suboscine split (∼50 Mya). This two-fold dynamism, characterized by both rapid genic turnover and ancient retention of genes, suggests that GRCs are constantly reshaped by gene gain, loss, retention and duplication ^1,18^. Moreover, the presence of W-derived genes on male GRCs hints at a role in hosting sexually antagonistic loci, echoing the GRC’s hypothesized role in mitigation of sexual conflict ^27^. The finding of W-derived genes in the House Finch GRC, along with the confirmation of extremely high abundances of repeats (see discussion below) refines our understanding of GRC composition, moving beyond earlier characterizations reporting lower repeat content ^1,4,5^.

### GRCs as permissive reservoirs for active ERVs

The accumulation and retention of intact, transcriptionally active ERVs strongly suggest that GRCs function as genomic reservoirs for these mobile elements. Unlike the A-chrs, where TE activity is often suppressed and insertions can be rapidly removed or silenced in avian species ^41^, GRCs appear to provide a more permissive environment for GRC retention and transposition. The GRC may act as a reservoir where TEs can persist without the stringent selective pressures present on the A-chrs, thereby shielding the somatic genome from harmful insertions while still permitting TE-driven variation to arise in the germ line. The significantly higher expression of ERV transcripts from the GRC compared to the A-chrs in House Finch germline-enriched tissue (Fig. 6) suggests that retrotransposons on the GRC are not merely relics of past insertions but instead remain transcriptionally active.

Two factors could contribute to the high ERV density and activity on the GRC. First, GRCs are eliminated during somatic cell development, potentially relaxing purifying selection against deleterious TE insertions that might otherwise affect organismal development or function. Our empirical findings therefore support the hypothesis that the GRC mitigates the deleterious effects of TEs in the avian soma (Vontzou et al. 2023). This pattern is also consistent with the observation of enriched TEs in programmed-eliminated DNA in other taxa, including nematodes, copepods and lamprey ^20^. Moreover, active transposons in a somatic genome are typically kept in check by host defense mechanisms through, for instance, DNA methylation ^46^ and piRNA pathways ^47^, and loss of such control leads to transposon reactivation, often with detrimental consequences ^8^. The elevated expression of ERVs on the GRC suggests a relaxation of these somatic repressive mechanisms on the GRC. One possibility is that the unique chromatin environment or regulatory landscape of the germline-specific GRC might also be less effective at silencing ERVs or facilitating their removal compared to the mechanisms operating on A-chrs. Overall, we interpret the avian GRC as an evolutionary reservoir: the host genome maintains its integrity and small size by expelling or compartmentalizing TEs onto the GRC, which in turn preserves those TEs in an environment where they can occasionally express and mobilize, seeding new insertions that natural selection can later harness as adaptive regulatory or coding novelties.

### Evolutionary implications for avian genome evolution and speciation

The temporal landscapes of ERV-LTR accumulation show a striking synchronicity: continuous ERV proliferation on GRCs begins around 50 Mya, at the base of passerine birds, and accelerates during the last ∼10 Ma, at the same point in time that Chen *et al.* ^17^ documented lineage-specific bursts of ERV solo-LTRs on A chrs. We propose that intact ERVs persist and amplify on GRCs, shielded from strong selection, and occasionally mobilize into A-chrs, where unequal recombination effectively converts most new insertions to solo LTRs, leaving a substrate and raw material for potential adaptive shifts while minimizing TE load. Indeed, transposition events in germ cells can induce mutations, chromosomal rearrangements, or novel regulatory sequences that may accelerate genetic divergence between populations (the “TE thrust” hypothesis; Oliver and Greene ^48,49^). For instance, the enrichment of specific classes of retroelements (ERV and L1 retroelements) causing cis-regulatory divergence between humans and chimpanzees has been deemed essential for the morphological changes to the cranium during primate evolution ^50,51^. Similarly, retrotransposons (ERVs) contribute to trophoblast gene expression driving placental evolution as enhancers or promoters ^52^. Thus the reservoir of intact ERVs on the GRC effectively increases the rate and quality of mutations for innovation in the A-chrs of the soma.

Concurrently, differential ERV activity can facilitate reproductive isolation, because active TEs are well-known to cause genomic conflicts in hybrids ^10,53^. Bursts of TE movement promote rearrangements via ectopic recombination, lowering hybrid fitness when divergent populations interbreed ^11,54^. A classic example is the P-element in Drosophila, where crosses between individuals with and without this transposon lead to hybrid dysgenesis and sterility ^55,56^. Similarly, diverging bird lineages with differing GRC-encoded ERVs or regulatory controls over them might produce hybrids with misregulated transposons, genomic instability, or defective germline development, thereby lowering hybrid fitness. Thus, GRC-permissive TE activity offers a plausible mechanism for accelerating post-zygotic barriers while simultaneously fostering adaptive divergence—the essence of the “TE-thrust” model ^48,49^

Although the correlations between GRC-mediated ERV bursts and speciation are compelling, causality remains inferential. Demographic processes, such as drift in small founder populations, could permit TE expansions independently of their role in driving speciation ^57^. It is also possible that GRCs and their transposons coevolve with speciation events in a feedback loop, rather than one simply causing the other. Future studies, for instance examining hybrids or experimental crosses, would be invaluable to test whether differences in GRC content and activity indeed result in incompatibilities or accelerated divergence. These insights extend our understanding of how TE dynamics can be compartmentalized and leveraged in genome evolution, highlighting the GRC as a peculiar but potentially powerful facilitator of avian diversification ^58,59^.

### Limitations and future directions

Despite the advancements offered by long-read sequencing, characterizing GRCs remains challenging. The assemblies presented here, although improving on earlier GRC assemblies, are still incomplete and fragmented, due to the repetitive nature of GRCs, which complicates assembly even with HiFi data. Furthermore, the subtractive approach used to identify GRC-specific sequences, although commonly used in GRC genomics, is inherently conservative and may omit GRC regions that retain high sequence similarity to A-chrs. Future research employing cytogenetics ^1^ or single-cell sequencing technologies or methods to physically isolate specific germline cell stages could provide more refined GRC sequences and cell-type-specific transcriptomic and epigenetic profiles. Additionally, investigating GRCs in females across different songbird lineages is crucial for a comprehensive understanding of their inheritance, function, and potential sex-specific roles in evolution.

In conclusion, our study positions the avian GRC as a hidden but potent driver of genome innovation—and, potentially, of avian biodiversity itself—by revealing its dynamic gene content, varied evolutionary origins, and exceptionally high repeat abundances, particularly of active ERVs. By housing these diverse elements, including active retroelements, in a chromosome that is excluded from the soma, birds appear to have evolved a clever compromise: protecting body cells from potential TE damage or gene misexpression while preserving a ‘sandbox’ where both TEs and duplicated genes can generate heritable novelty.

## Methods

### Sampling and sequencing

A previously published pangenome study ^23^ provided a House Finch heart (somatic) assembly and a Common Rosefinch testis (germline) assembly, both generated with PacBio HiFi (highly accurate long-read) data. To assemble both GRCs and A-chrs, we obtained additional high-depth PacBio HiFi data from the same male individuals accessioned at the Museum of Comparative Zoology (MCZ), Harvard University (Table S1). Specifically, we sequenced a new House Finch testis sample at 53.4× coverage, a new Common Rosefinch heart sample at 45.6× coverage, and increased the Common Rosefinch testis coverage from 37.4× to 98.5×.

We extracted high-molecular-weight (HMW) DNA from these samples using the Qiagen MagAttract HMW DNA Kit. DNA quality was checked using TapeStation (Agilent), Qubit, and NanoDrop (Thermo Fisher Scientific) at the Bauer Core Facility at Harvard University, where HiFi sequencing was performed. PacBio HiFi libraries were prepared with the SMRTbell Express Template Prep Kit 2.0 (PacBio) and size-selected on a BluePippin system (Sage Science), then sequenced on Sequel IIe SMRT Cells (PacBio). To identify House Finch GRC sequences (see below), we performed whole-genome sequencing (WGS) on testis (germline) and heart (somatic) tissues from two individuals. DNA was extracted using the DNeasy Blood and Tissue Kit (Qiagen) according to the manufacturer’s protocol. Libraries were prepared using transposase-mediated fragmentation and sequenced on an Illumina NovaSeq S4 platform (2×150 bp) at the Bauer Core Facility.

For RNA sequencing, we sampled three tissues (testis, brain, heart) from four House Finch individuals for GRC gene annotation and transcriptomic analyses, and three tissues (testis, muscle, heart) from one Common Rosefinch individual for its A-chr and GRC gene annotation. Total RNA was extracted with the RNeasy Mini Kit (Qiagen), and its quality was evaluated with a TapeStation (Agilent). Libraries were prepared using the KAPA mRNA HyperPrep Kit (Roche). After final quality checks (TapeStation and qPCR), libraries were pooled and sequenced on an Illumina NovaSeq S4 platform (2×150 bp) at the Bauer Core Facility. The total number of raw reads, overall alignment rate, and average exonic coverage for each sample are listed in SI Appendix, Table S1, which also contains detailed information about each sample (including sex, locality [GPS], and sequencing depth).

### GRC and A-chr Assembly

We aimed to assemble GRC contigs from sequences that are divergent from A-chrs, following approaches similar to previous GRC genomic studies ^21,25^. Because there is no standardized pipeline due to varying sequencing strategies, we adopted a four-step process using PacBio HiFi data: (1) mapping somatic reads to the testis genome (combined genome: A-chrs plus GRC); (2) identifying GRC-specific regions; (3) extracting GRC reads mapping to GRC-specific regions; and (4) assembling GRC assemblies from GRC reads. Specifically, we first generated somatic and germline assemblies for both species from the HiFi data (see above). PacBio HiFi BAM files were converted to FASTQ using bam2fastx (PacBio), and Hifiasm was used for assembly (Cheng et al. 2021). Assembly quality was assessed using assembly-stats (https://github.com/sanger-pathogens/assembly-stats) (SI Appendix, Table 1). Second, we identified GRC-specific regions by mapping somatic HiFi reads to the germline assembly (minimap2 with the -x map-hifi preset). We calculated mean mapping depth per 100 bp window using samtools and bedtools, designating windows with zero coverage as GRC-specific regions. Third, we extracted GRC reads by mapping germline HiFi reads back to the germline assembly using Hifiasm, then isolating those reads mapping to GRC-specific regions with bedtools and bioawk ^60^. Fourth, we assembled these extracted GRC reads with Hifiasm to produce final GRC contigs.

Because more genomic resources were available for House Finch (including HiFi and WGS from Fang and Edwards ^23^), we performed extra analyses to identify high-confidence GRC reads. In Step (2), we incorporated somatic HiFi reads from eight additional male individuals from Fang and Edwards ^23^(total ∼184× coverage) for more stringent identification of diverging regions between GRC and A-chrs. Next, for GRC sequences similar or indistinguishable from A-chr sequences, we searched for germline-specific SNPs across three individuals (one HiFi and two WGS). We mapped both somatic and germline WGS reads to the HiFi somatic assembly using minimap2 in the preset for mapping short paired-end reads (-ax sr). We then identified SNPs differentiating germline from somatic genotype calls (where the genotype of the somatic sample is ‘0/0’ and that of the germline is ‘0/1’ or ‘1/1’ in the same individual) across all three individuals using bcftools and vcftools ^61,62^. Any germline HiFi reads containing these SNPs were extracted as GRC reads. Finally, these GRC reads (both divergent and similar to A-chrs) were combined and assembled to generate GRC contigs for House Finch.

To validate the assembled GRCs, we mapped somatic and germline HiFi reads to the combined genome (A-chr plus GRC) in both species using minimap2. Median read depth per 100 bp window was calculated with samtools ^62^ and bedtools. The GRC assemblies were confirmed by: (1) the GRC contigs showed zero coverage from somatic reads, indicating that GRC contigs were unique to germline tissue; (2) in testis, GRC coverage was lower than A-chr coverage (House Finch: 102.32 vs. 21.98; Common Rosefinch: 99.77 vs. 29.35), consistent with the GRC’s haploid status and presence only in spermatocytes (as observed in the Blue Tit GRC study); and (3) the GRC:A-chr coverage ratios (House Finch = 0.22, Common Rosefinch = 0.29) closely matched ratios reported in other songbird testis-based GRC sequencing studies ^5,21,25^.

### Gene Annotation

The coding genes of the House Finch GRC and the Common Rosefinch GRC and A-chrs were annotated using (1) RNA-seq data, (2) protein-based homology, and (3) ab initio gene prediction. Specifically, RNA sequencing reads from both somatic and germline tissues were quality-checked with FASTQC ^63^ and trimmed with Trim Galore ^64^ to remove low-quality bases and adapters, then mapped to the respective assemblies using hisat2 (Kim et al. 2019). Transcripts were constructed per tissue using StringTie ^65^, and open reading frames (ORFs) were predicted with TransDecoder (<u>github.com/TransDecoder/</u>). We also employed Galba ^66^, which combines miniprot ^67^, a protein-to-genome aligner, and AUGUSTUS ^68^, an ab initio gene predictor, using protein sequences of House Finch, chicken, and zebra finch retrieved from NCBI. All gene models generated by these approaches were integrated with StringTie-merge, and duplicates or incomplete structures were removed using AGAT (<u>github.com/NBISweden/AGAT</u>). Gene IDs in the final GFF were matched with NCBI gene IDs using blastx ^26^. We conducted Gene Ontology (GO) enrichment analysis for GRC-linked genes with clusterProfiler (enrichGO) ^69^ and visualized the results as dot plots.

### Repeat and Transposable Element (TE) Analysis

We performed repeat annotation of the GRC and A-chr assemblies for House Finch, Common Rosefinch, and Blue Tit using the RepeatModeler and RepeatMasker pipeline ^30^. First, de novo prediction of repetitive elements was carried out with RepeatModeler, which incorporates RECON ^30^ , RepeatScout ^70^, and Tandem Repeats Finder ^71^. We then concatenated these de novo libraries with the avian Repbase transposable element library ^72^ to form a curated repeat library, which served as input for RepeatMasker to annotate repetitive elements in each assembly. To accurately identify long terminal repeat retrotransposons (LTR-RTs), we used LTR_retriever ^31^, which integrates outputs from LTR_harvest ^32^ and LTR_FINDER ^33^ to produce high-quality, non-redundant LTR libraries and annotations. The resulting LTR library was again used by RepeatMasker for final LTR annotation. Solo-LTRs were identified using the “solo_finder.pl” script (LTR_retriever package) under the following criteria: (i) ≥80% coverage of the intact LTR entry, (ii) alignment score >300, (iii) length ≥100 bp, and (iv) no internal LTR region within 300 bp. Finally, we applied the “solo_intact_ratio.pl” script (LTR_retriever) to calculate the ratio of solo to intact LTRs for each family. The TE landscapes were obtained from the RepeatMasker outputs using a Perl script (https://github.com/4ureliek/Parsing-RepeatMasker-Outputs/blob/master/parseRM.pl) with an input of a substitution rate of 3.3×10⁻³ substitutions per site per million years for Passeriformes (referencing Zhang et al. ^73^).

### Phylogenetic analyses of GRC genes

To reconstruct the evolutionary history of the GRC, 12 GRC-linked genes shared among at least three bird species out of five published GRCs were used for phylogenetic analysis. Coding sequences (CDS) of these target genes were retrieved using ncbi_datasets for Chicken (*Gallus gallus*), Rifleman (*Acanthisitta chloris*), and Suboscines species, Blue-capped Manakin (*Lepidothrix coronata*), and Golden-collared Manakin (*Manacus vitellinus*). Multiple sequence alignments were performed using MAFFT ^74^ with the --localpair and --maxiterate 1,000 options, along with -- adjustdirectionaccurately to handle potential strand reversals. Alignments were processed with ClipKIT ^75^ to remove poorly aligned or phylogenetically uninformative sites. Phylogenetic trees were constructed using IQ-TREE ^76^ with 1,000 ultrafast bootstrap replicates (-B 1000). To assess the robustness of inferred topologies, we performed an approximately unbiased (AU) topology test in IQ-TREE with 1,000 bootstrap replicates (-zb 1000).

### Transcriptomic analysis

After quality control (see Gene Annotation), somatic RNA-seq reads were mapped to the somatic (A-chr) assembly, and gonadal RNA-seq reads were mapped to the germline (A-chr plus GRC) assembly. Alignments were then filtered using the featureCounts function (SUBREAD package) (Liao et al. 2013) in paired-end mode, retaining only uniquely mapped reads. To quantify expression of endogenous retroviruses (ERVs), we used featureCounts with genomic intervals corresponding to the protein-coding regions of annotated complete LTR retrotransposons (LTR-RTs). Per-genome counts were derived from the read counts and lengths of the corresponding ERVs. For differential expression analysis, we employed DESeq2 ^77^, reporting normalized expression in transcripts per million (TPM) adjusted for sequencing depth and gene length. ERV expression was compared between somatic and gonadal tissues, and between A-chrs and GRC in gonadal tissue. Because the proportion of germ cells in gonadal samples was uncertain, normalization was repeated assuming 10%, 5%, and 1% germ cell composition, and all results are shown (Fig. 6). Statistical significance was assessed using the Wald test, with p-values adjusted (padj) for multiple testing.

### Resource availability

The assemblies, raw HiFi, whole-genome sequencing and RNA data will be made accessible from the NCBI (PRJNAxxxx). Analytical scripts are available on GitHub at https://github.com/fangbohao/Germline_Restricted_Chromosome_Project.

## Acknowledgments

We thank Alexandria DiGiacomo and Kelsie Lopez for technical advice in RNA isolation and analysis. We also thank Jonathan Schmitt for contributions to the House Finch and Rosefinch collections at the Museum of Comparative Zoology, Harvard University. We acknowledge the FASRC Cannon cluster at Harvard University for computational resources. We thank the reviewers for their constructive comments. This work was funded by Harvard University, the Harvard Global Institute, and Harvard China Funds (to S.V.E. and B.F.).

## Author contributions

B.F. and S.V.E. designed research; B.F. performed research, analyzed data, and wrote original draft; S.V.E. supervised and funded project. All authors read and approved the bioRxiv manuscript.

