## Supplementary Material for "Avian germline-restricted chromosomes are reservoirs for active long-terminal-repeat retroviruses"

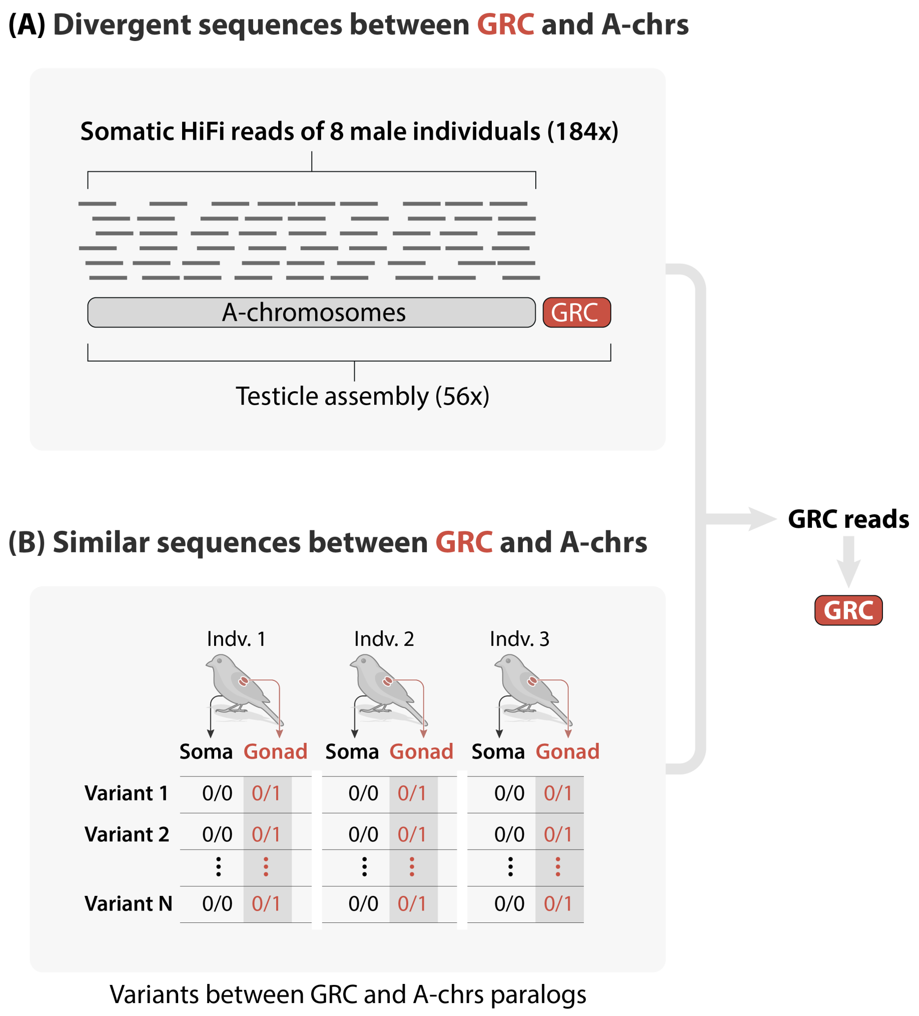


**Figure S1. Strategy for isolating germline-restricted chromosome (GRC) reads for *de novo* assembly (related to STAR Methods, "GRC and A-chr Assembly"). (A)** Identifying highly divergent GRC regions. PacBio HiFi reads from somatic tissues of eight male House Finches (total 184× coverage) were mapped to the testis HiFi assembly (56×). Genomic intervals showing zero somatic coverage (red) were classified as GRC-specific, whereas regions with coverage were retained as A-chrs (grey). Germline reads mapping to the GRC-specific intervals were extracted for downstream assembly. **(B)** Capturing GRC segments that remain similar to A-chr paralogs. For three additional males of House finches, short-read whole-genome data from soma and gonad were jointly genotyped. Sites that were homozygous reference in soma (0/0) but heterozygous or homozygous alternate in gonad (0/1 or 1/1; highlighted in red) were treated as germline-specific single-nucleotide polymorphisms. Only SNPs shared all three males are deemed as high-confidence SNPs. HiFi reads containing these SNPs were retrieved, allowing recovery of GRC sequences that are otherwise indistinguishable from their A-chr paralogs.

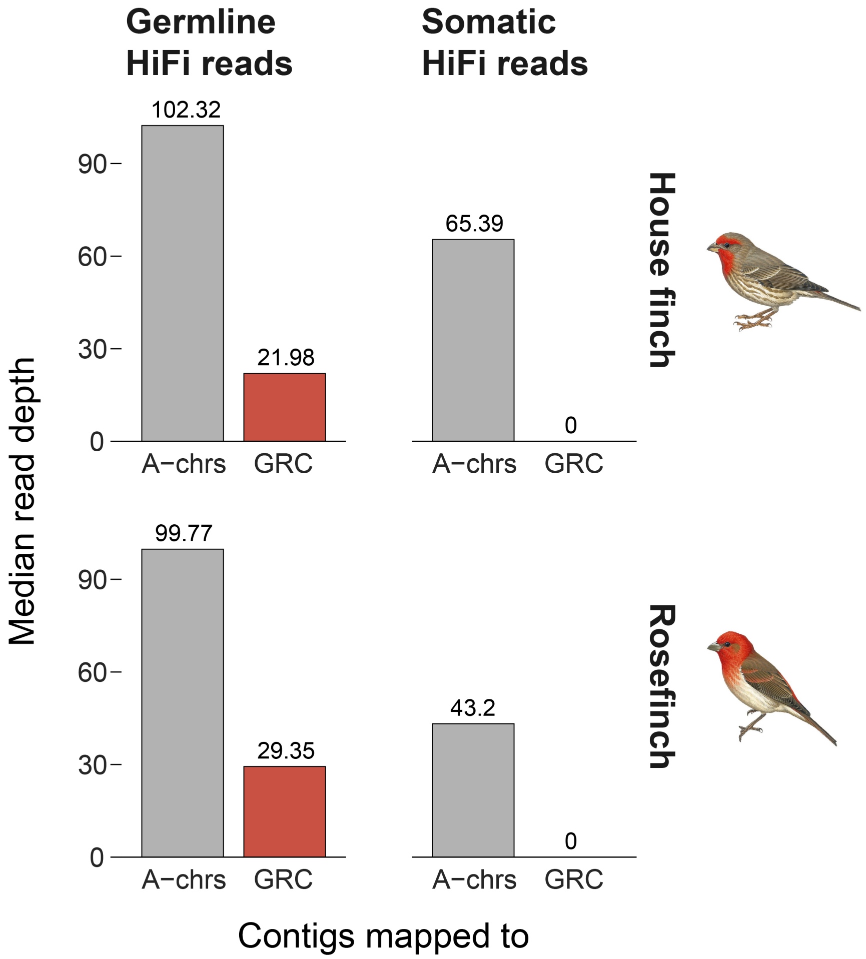


**Figure S2. Validation of House Finch (HF) and Common Rosefinch (RF) GRC assemblies using read coverage.** HiFi read mapping coverage from somatic (heart) and germline (testis) tissues onto the combined A-chrs and GRC assemblies. GRC contigs show near-zero somatic coverage, confirming germline restriction. Lower coverage in testis for GRCs compared to A-chrs (mean GRC/A-chr ratios: HF ≈0.22, RF ≈0.29) reflects their haploid status and presence only in specific germline cells.

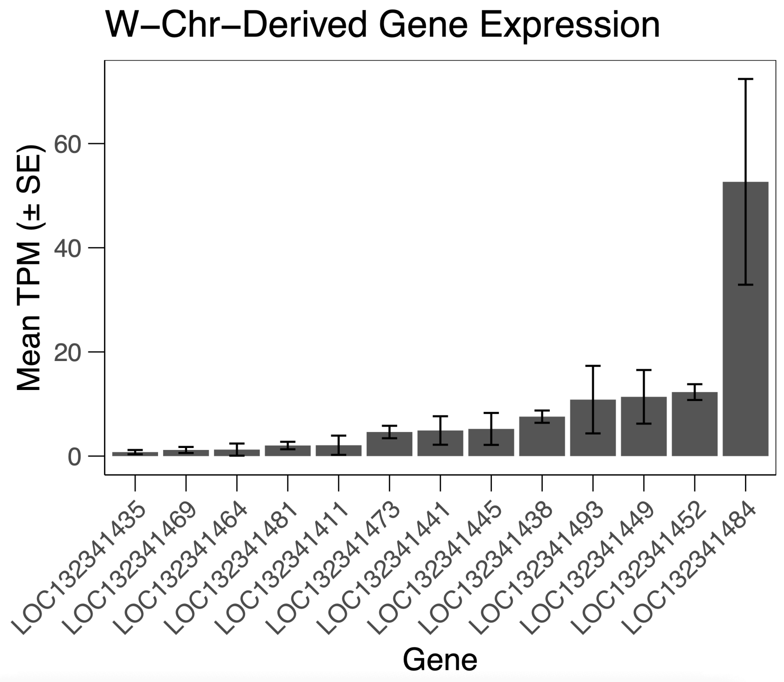


**Figure S3. Transcriptional activity of W-chromosome–derived paralogs located on the male House Finch GRC (related to Figure 1 and STAR Methods, "Gene Annotation" and "Transcriptomic Analysis").** Bar plot shows the mean transcripts-per-million (TPM ± s.e.) for the 13 GRC-linked genes whose closest paralogs reside on the W chromosome. Expression was quantified from testis RNA-seq data of three adult males (biological replicates) using *featureCounts* followed by *DESeq2* length-normalization (Methods). At least eight loci exhibit detectable germline expression, with LOC132341484 displaying the highest activity (~50 TPM). All W-derived genes are shown.

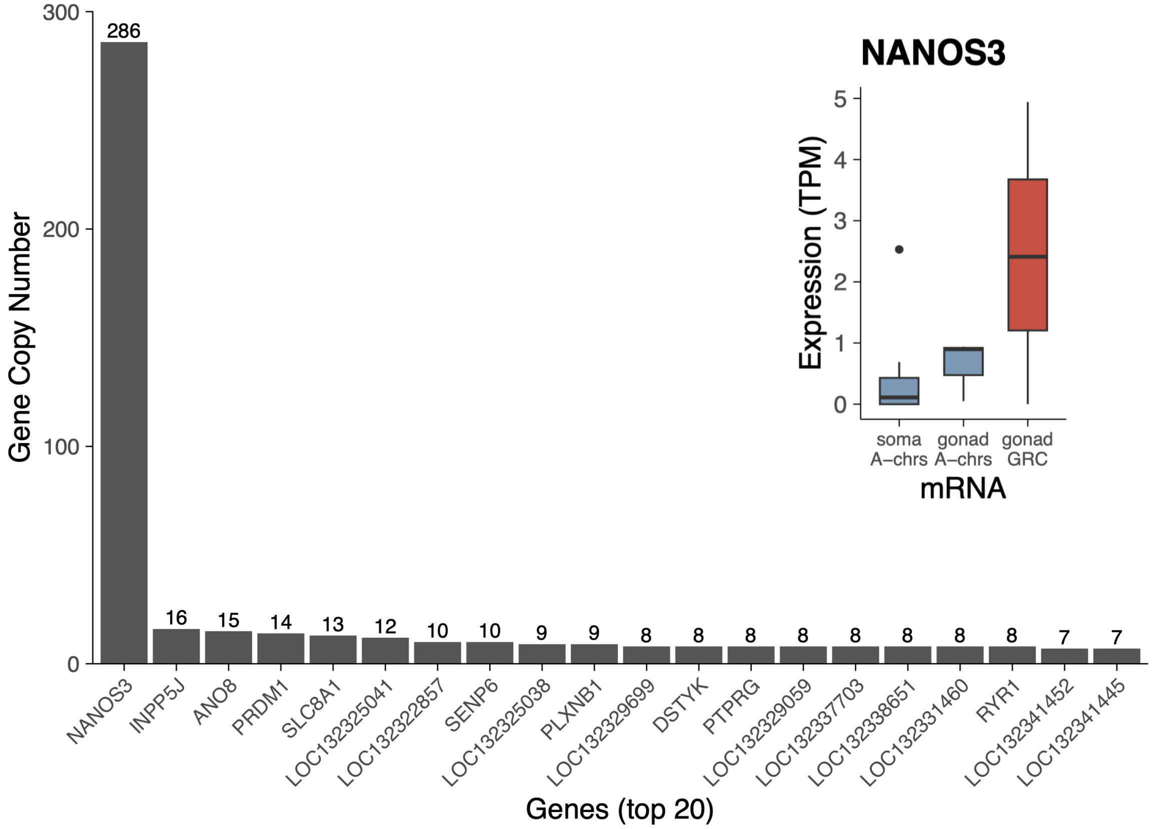


**Figure S3. Gene duplications on House Finch GRC.** Left, copy number for the 20 most duplicated GRC genes; NANOS3 is expanded to 286 copies and just one functional (with transcription), far exceeding any other locus. Right inset, NANOS3 mRNA levels (TPM) of the expressed gene across three males: negligible in somatic tissues, low from A-chromosomes in testis, and highest from the GRC, indicating germline-restricted transcription.

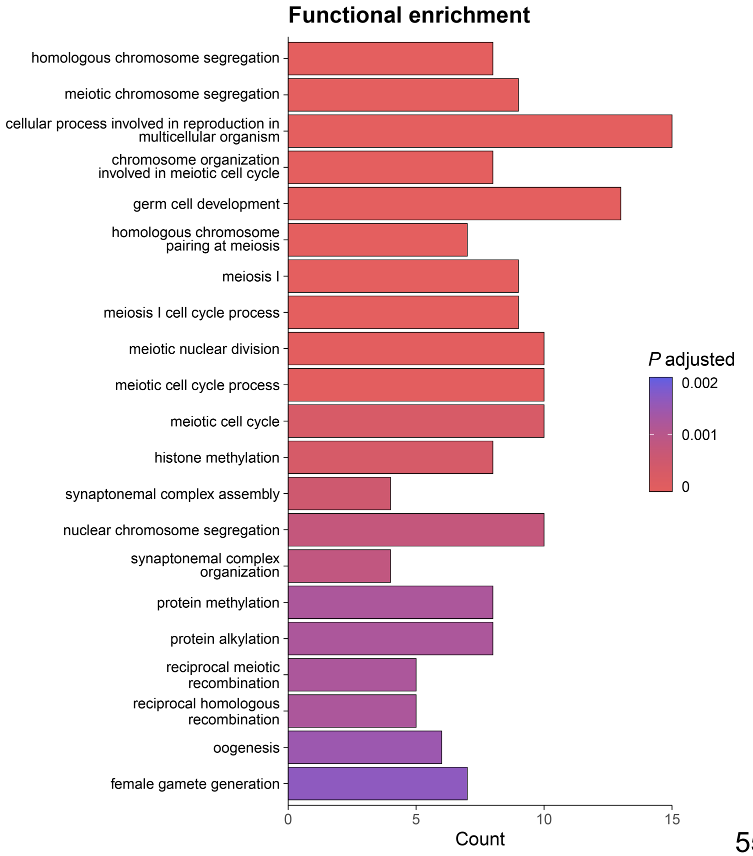




**Fig. S5. Gene Ontology (GO) enrichment analysis of GRC-linked genes.** Dot plot displaying significantly enriched GO terms associated with GRC-linked genes across avian species. C Over-represented categories center on meiosis, chromosome segregation, germ-cell development, and associated cell-cycle processes, highlighting the germline-biased function of the GRC gene set. Axes typically represent GO terms and significance (-log10(p-value)) or gene ratios, with dot size/color indicating gene counts or enrichment scores.

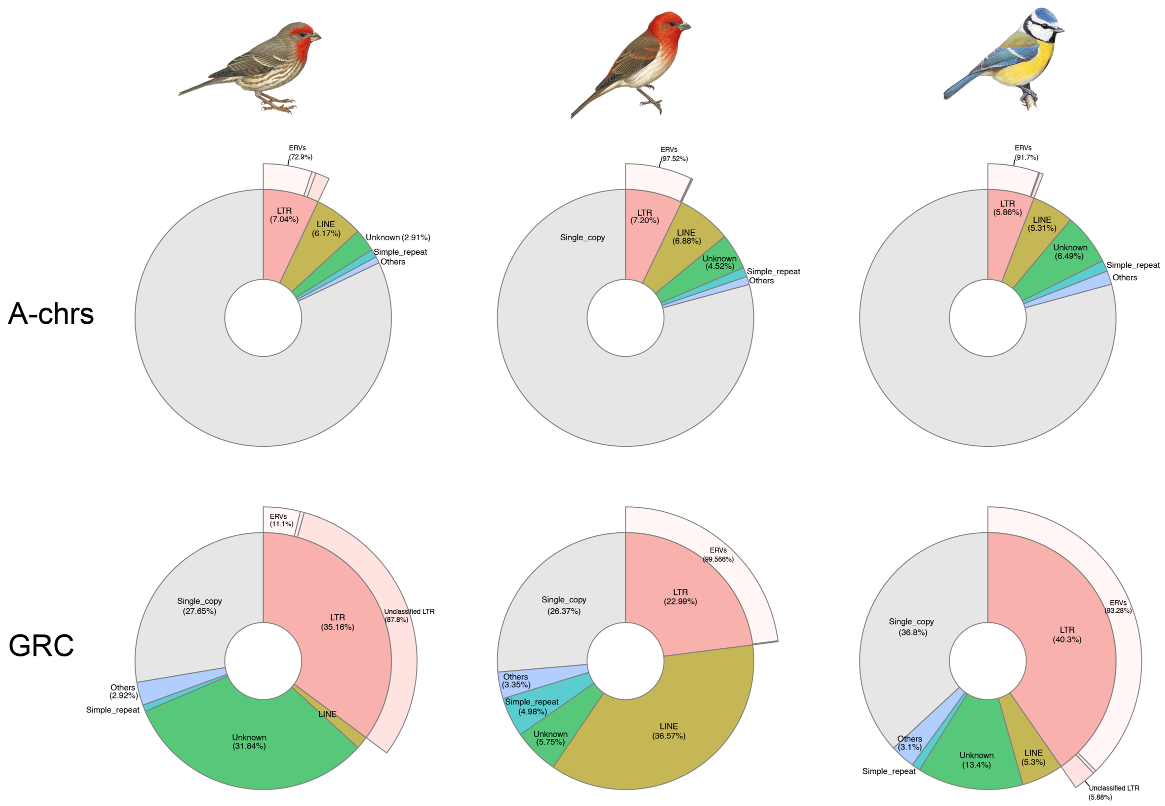


**Figure S5. Composition of major repeat classes in A-chrs and GRCs.**
Donut charts show the percentage of each repeat class in A-chrs (top row) and GRCs (bottom row) of House Finch (left), Common Rosefinch (center) and Blue Tit (right). Segments are color-coded and labelled with their fractional contribution. This figure complements Figure 3 by showing the dominance of ERVs within the LTR category. Values correspond to the summaries in Table S4.

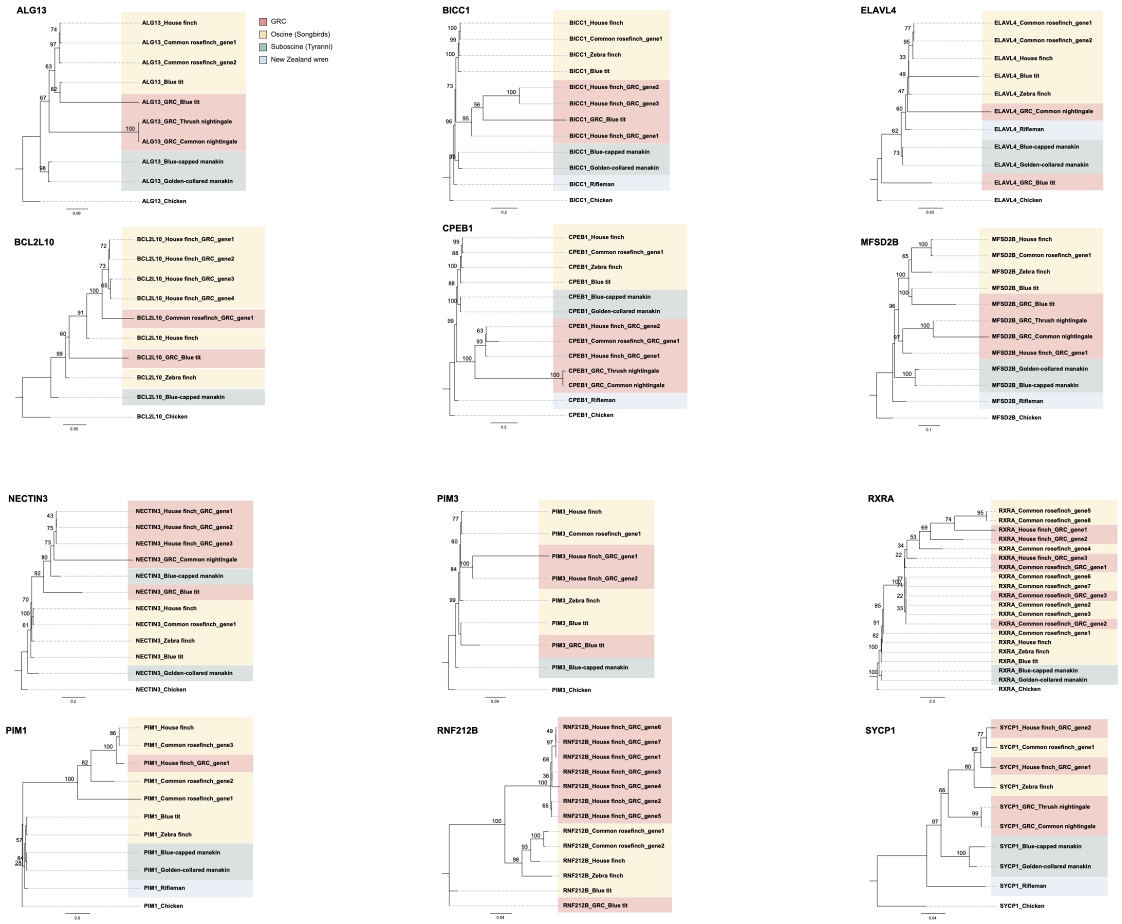


**Figure S6 | Phylogenies of the 12 GRC-linked genes shared by ≥ 3 passerines.**
Maximum-likelihood trees (IQ-TREE, 1,000 ultrafast bootstraps) are shown for each gene.
